# Non-photosynthetic bacteria produce photocurrent mediated by NADH

**DOI:** 10.1101/2023.01.16.524302

**Authors:** Yaniv Shlosberg, Jakkarin Limwongyut, Alex S. Moreland, Guillermo C. Bazan

**Affiliations:** Department of Chemistry and Biochemistry, University of California at Santa Barbara, Santa Barbara, CA, 93106, United States

## Abstract

In recent years, the concern from the global climate change has driven an urgent need to develop clean energy technologies that do not involve combustion process that emit carbon into the atmosphere. A promising concept is microbial fuel cells that utilize bacteria as electron donors in a bio-electrochemical cell performing a direct electron transfer via conductive protein complexes or by secretion of redox active metabolites such as quinone or phenazine derivatives. In the case of photosynthetic bacteria (cyanobacteria) electrons can also be extracted from the photosynthetic pathway mediated mostly by NADH and NADPH. In this work, we show for the first time that the intact non-photosynthetic bacteria *Escherichia coli* can produce photocurrent that is enhanced upon addition of an exogenous electron mediator. Furthermore, we apply 2D-fluorescence measurement to show that NADH is released from the bacterial cells, which may apply as a native electron mediator in microbial fuel cells.

## Introduction

In recent years, energy innovations are directed toward the development of clean technologies that do not involve combustion processes and in this way can reduce carbon emission. A promising energy solution that is extensively developed is microbial fuel cells (MFCs)^1^. The main approach of MFCs is to exploit the ability of bacterial cells to donate electrons at the anode^2,3^ of a bio-electrochemical cell. Alternatively, several species can also apply as electron acceptors at the cathode^2,4,5^. The bacterial cells perform external electron transport by 2 kinds of mechanisms: direct and mediated electron transfer. Direct electron transfer (DET) is conducted by metal respiratory (MTR) complexes in the cellular membrane^6–9^. Mediated electron transfer is performed by native secretion of redox active molecules. Among these molecules are various derivatives of quinones and phenazines ^10–15^. The electrical current production can be enhanced by addition of artificial exogenous electron mediators such as cystine, neutral red, thionin, sulphides, ferric chelated complexes, quinones, phenazines, and humic acids^16–21^. Among the most efficient bacterial species in MFCs are *Shewanella oneidensis* and *Geobacter sulfurreducens* which consist of a relatively high amount of *pili* and cytochrome types that are capable of charge transfer^6–9,22,23^. MFCs are not limited to bacteria and can also utilize yeasts^24^. A unique class of MFCs is the bio-photoelectrochemical cells (BPECs)^25–30^. This approach utilize the ability of photosynthetic organisms to perform external electron transfer, while the source of the electrons originate from both the respiratory and photosynthetic pathways^31^. The major electron mediators in BPECs are NADH and NADPH that can cycle electrons from photosystem I inside the cells and the external anode of the BPEC^25^. Enhancement of the photocurrent can be achieved by the exogenously adding natural electron mediators such as NADH, NADPH and vitamin b1 or the non-natural mediator potassium ferricyanide^30^.

In this work, we show for the first time that it is possible to produce photocurrent from non-photosynthetic bacteria in a BPEC while the electron transfer is being mediated by NADH.

## Results and Discussion

### *E.coli releases* NADH and FAD to the external cellular medium

Recent works have reported the release of the redox active molecules Nicotinamide adenine dinucleotide (NADH) and Nicotinamide adenine dinucleotide phosphate (NADPH) from various organisms such as cyanobacteria^25,30^, microalgae^26^, seaweeds^32^, plant’s leaves^29,34,36^, roots^38^ and sea anemones^39^. These molecules can apply as electron mediators in bio-electrochemical cells catalysing electrons transport between the respiration and photosynthetic pathways of the cells and the external anode. The biological reason for the release of NADH and NADPH is not fully understood. It was suggested that it may derive from a minor leak of cytoplasmatic content, be involved in quorum sensing or apply to reduce iron to internalise it^25^. Mediated electron transport in non-photosynthetic bacteria in microbial fuel cells (MFCs) was previously explained by secretion of quinone and phenazines derivatives^16–21^. Nevertheless, the possibility of a release of NADH and NADPH from non-photosynthetic bacteria was not studied yet. To investigate this, we wished to analyse the external cellular solution of *E.coli* cells by 2D – fluorescence measurements. *E.coli* cells were were centrifuged (20 min, 6000 rpm) and resuspended in phosphate buffer saline (PBS) After 1 h of incubation, the cells were filtered by 0.22µm filter. 2D-fluorescence of the filtrate was measured (λ_ex_ = 260 – 500 nm, λ_em_ = 280 – 580 nm) (Fig. 1). The fluorescence maps showed peaks with maxima around (λ_ex_ **=** 280 nm, λ_ex_ **=** 310 nm), (λ_ex_ **=** 280 nm, λ_ex_ **=** 350 nm), (λ_ex_ **=** 300 nm, λ_ex_ **=** 450 nm), (λ_ex_ **=** 360 nm, λ_ex_ **=** 450 nm), and (λ_ex_ **=** 440 nm, λ_ex_ **=** 520 nm) that are correlated with the spectral fingerprints of tyrosine, tryptophane, NAD^+^ or NADP^+^, NADH or NADPH, and FAD ^25,33,35^. As *E.coli* not a photoautotroph and cannot produce NADPH in a photosynthetic pathway like cyanobacteria, we suggest that the peaks with maxima at (λ_ex_ **=** 300 nm, λ_ex_ **=** 450 nm) and (λ_ex_ **=** 360 nm, λ_ex_ **=** 450 nm) originate from the fluorescence of NAD^+^ and NADH respectively. Based on the obtained results, we suggest that the identified redox active species NADH and FAD may apply as native electron mediators in MFCs in addition to quinones and phenazines.

**Fig. 1.**
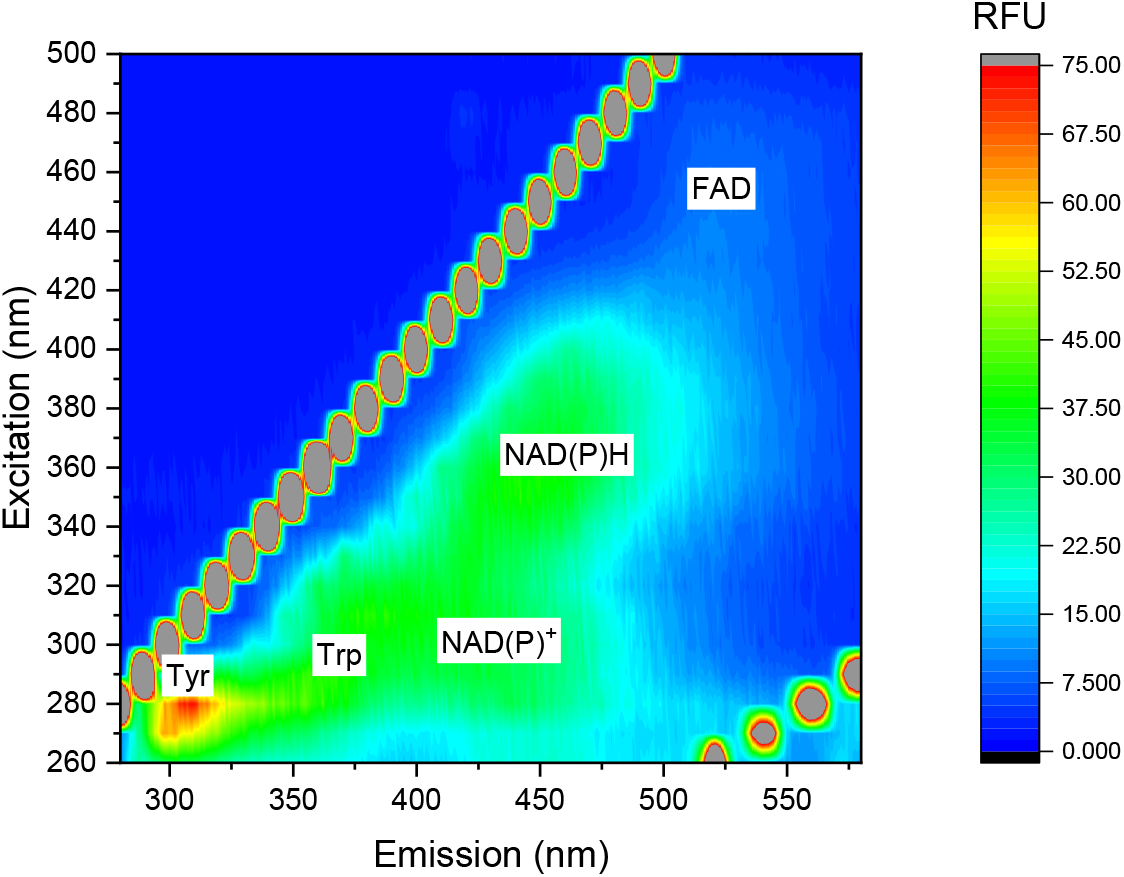
*E.coli* releases NADH and FAD to the external cellular medium. 2D-fluorescence spectra of the external solution of *E.coli* was measured. White labels show the maxima of the peaks of tyrosine, tryptophan, NAD^+^, NADH, and FAD. The lines of diagonal spots that appear in all of the maps presented here and in the following figures results from light scattering of the Xenon lamp^37^.

### Light irradiation of *E.coli* enhances the release of electron donors

Previous works have reported the ability of photosynthetic bacteria (cyanobacteria) to produce photocurrent in bio-photo electrochemical cells. The major source of electrons in such systems is the photosynthetic pathway to produce NADPH molecules. A small portion of NADPH is also released from the cyanobacterial cells into the external solution. This release is enhanced upon association of the cells with the anode of an electrochemical cell. Another molecule that significantly contributes to the current production in cyanobacteria is NADH that is formed in the tricarboxylic acid cycle (TCA cycle). Recent studies have shown that light irradiation on *E.coli* cells can enhance the reduction of NAD^+^ to NADH in the TCA cycle^40^. Based on this and the identification of NADH in the external solution (Fig. 1), we sought irradiation of *E.coli* that may enhance the current production by *E.coli* when applied as electron donor in MFCs. To explore this, we have designed a bio-electrochemical system that consist of screen-printed electrodes with a graphite working electrode (WE), graphite counter electrode (CE), and a Silver coated with silver chloride reference electrode (RE). A drop of 100 µL of *E.coli* suspension (10^6^ CFU / mL) was placed on the screen-printed electrode. Light irradiation was done from top using a white LED. (Fig. 2a,b)The light intensity at the height of the electrode surface was 1 Sun (1000W / m^2^). Cyclic voltammetry of was measured in dark and light at potential varied from 0 to 1 V, with a scan rate of 0.1 V/s. The voltammogram showed 2 peaks at maximal potentials of 0.6 and 0.9 V that were enhanced under illumination (Fig. 2c). The bigger peak around 0.9 V correlates with the voltammogramic fingerprint of NADH ^41^, strengthening our hypothesis that this molecules plays a key role in the electron mediation between the bacterial cells and the anode.

**Fig. 2.**
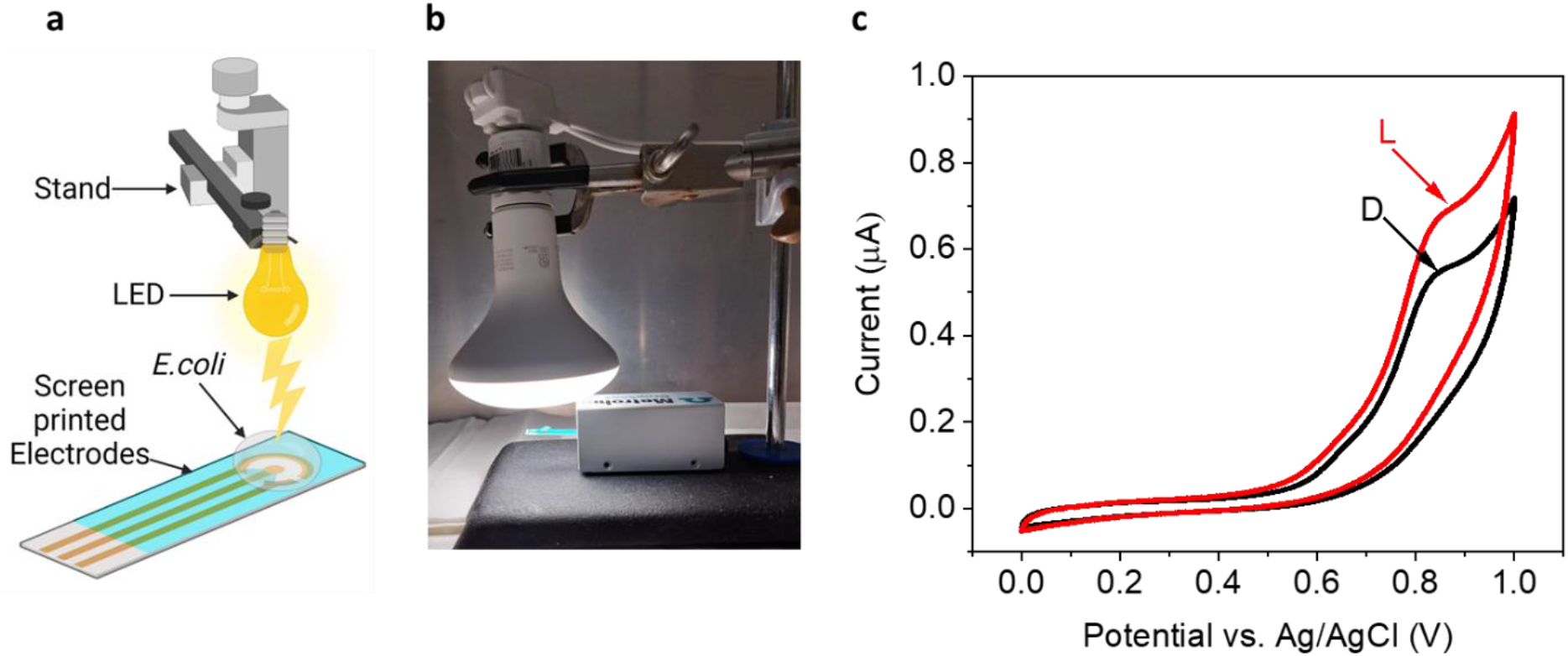
Light irradiation of *E.coli* enhances the release of electron donors. Cyclic voltammetry of *E.coli* in dark and light. **a** A schematic illustration of the system that consist of a white Ledirradiating from top, screen-printed electrode, and a drop of *E.coli*. **b** A picture of the system. **c** Cylclic voltammetry measurements of the bacterial cells in dark (black) and light (red).

### *E.coli* produces photocurrent in a bio-electrochemical cell

Next, we wished to explore whether *E.coli* can produce photocurrent in a bio-electrochemical cell. Chronoamperometry of *E.coli* was measured with dark/light irradiation intervals of 100 s. A potential bias of 0.9 V was applied to the WE (Fig. 3). This potential was chosen based on the bigger peak that was obtained in the cyclic voltammetry measurements (Fig. 2). A current of about 0.07 µA /cm^2^ was obtained in dark. Upon irradiation, the current was enhanced by ∼ 0.05 µA /cm^2^. PBS applied as a control, showing no significant dark current and a photocurrent of ∼ 0.005 µA /cm^2^ that was obtained because of a direct light absorption in the WE. We postulated that this effect was significantly smaller for the bacterial suspension as their turbidity do not allow the same transparency as the clear PBS solution. Previous studies about photocurrent production from various intact cyanobacterial species, showed a photocurrent of about 0.3 µA /cm^2^ (without being normalised to their chlorophyll content), while about 25 % of the current (∼ 0.075 µA /cm^2^) was reported to originate from NADH. A similar photocurrent production was obtained for *E.coli* (Fig. 3). Based on this, we suggest that in cyanobacterial based BPECs part of the photocurrent does not originate from photosynthesis but from and enhanced formation of NADH catalysed by light irradiation.

**Fig. 3.**
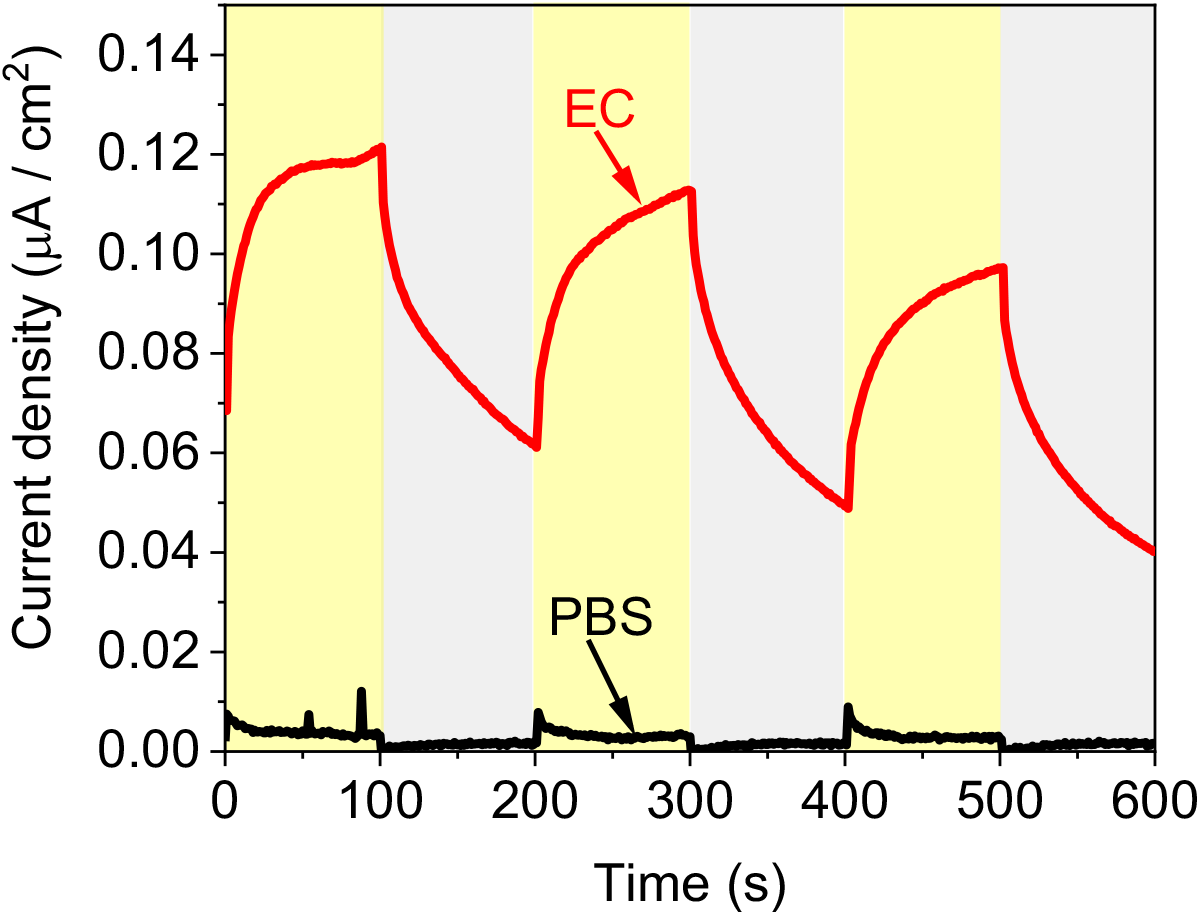
*E.coli* produces photocurrent in a bio-electrochemical cell. Chronoamperometry measurements of PBS (black) and *E.coli* (red) and were conducted under dark/light illumination intervals with an applied potential of 0.9 V on the anode. Gray and yellow coluors represent partial times of the measurement that were conducted in dark and light respectively.

### The electron transfer mechanism in photo microbial fuel cells

Based on the results obtained in this study, and previous studies we suggest mechanisms for the external transfer between the bacterial cells and the anode. In dark and upon association of the bacteria with the active electrochemical cells. NADH molecules are released from the cells donating electrons at the anode to produce electrical current. The oxidized form NAD^+^ is internalized into the cells cytoplasm were it may be re-reduced to NADH in the Glycolysis pathway. Upon irradiation, the reduction of NAD^+^ to NADH is being enhanced, increasing the number of molecules in the NADH cytoplasmatic pools, and the release of these molecules from the bacterial cells to produce electrical current. FADH_2_ may also be released from the cells to be oxidized at the anode into FAD, that can be reduced again by MTR complexes. These MTR complexes can also conduct a DET to donate electrons at the anode. DET may also be performed by conductive *pili*. Some of the non-light dependant electrical current may also derive from secretion of quinone and Phenazine molecules.

**Fig. 4.**
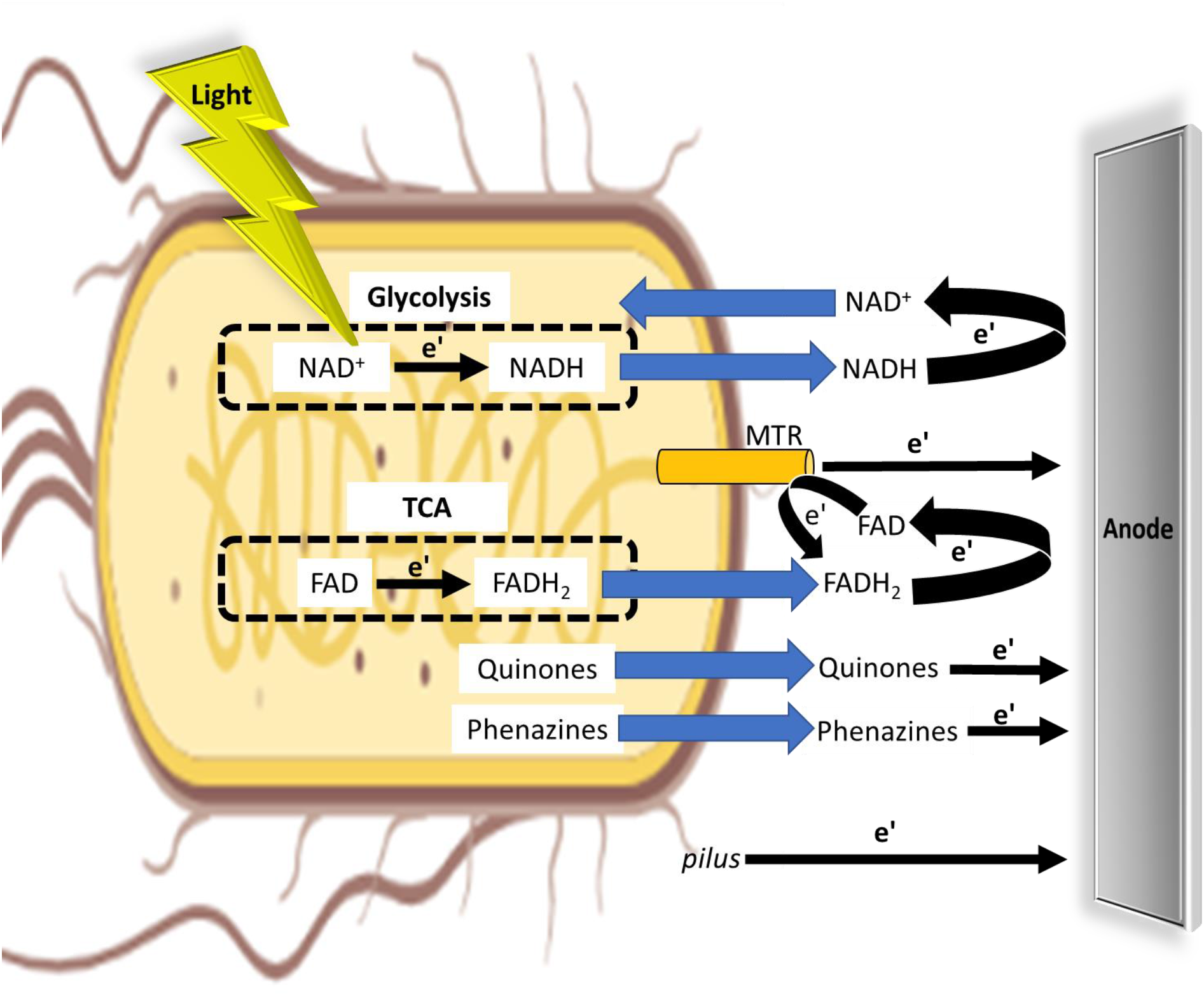
The electron transfer mechanism in photo microbial fuel cells. A schematic description of the different light dependent and independent external electron transfer mechanisms in MFCs. NAD^+^ is being reduced to NADH in the Glycolysis pathway. This reaction is induced under light irradiation. FAD is being reduced to form FADH_2_ in the TCA cycle. Upon association of the bacteria with the active bio-electrochemical cell, NADH, FADH_2_, Quinones and Phenazines are released from the bacteria to donate electrons at the anode. NAD^+^ molecules can re-enter the bacterial cell to be re-reduced. FAD may be re-reduced by MTR complexes. Some of the electrical current may also originate from a direct electron transfer between MTR complexes or *pili* and the anode. Dashed circles represent the Glycolysis and TCA cycles. Blue arrows represent the trafficking of molecules inside and outside of the bacteria. Black arrows represent electron transfer.

## Conclusions

In this work, we showed that non-photosynthetic bacteria can produce photocurrent. We applied 2D-fluorescence spectra of the external solution and cyclic voltammetry of the bacterial cells to show for the first time that some of the electrical current generation originate from NADH. These discoveries may help to improve the current production in MFCs.

## Acknowledgments

This work was supported by the National Institutes of Health R35 Maximizing Investigators’ Research Award (MIRA, R35GM142920). Yaniv Shlosberg is funded by the Otis Williams fellowship.

Some of the figures were prepared using Biorender.com

## Author contributions

YS conceived the idea. YS designed the experiments. YS performed the main experiments. JL and AM assisted performing some of the experiments. YS, JL, GB and LS wrote the manuscript. YS and LS supervised the entire research project.

